# Permutation Testing in the Presence of Polygenic Variation

**DOI:** 10.1101/014571

**Authors:** Mark Abney

## Abstract

This article discusses problems with and solutions to performing valid permutation tests for quantitative trait loci in the presence of polygenic effects. Although permutation testing is a popular approach for determining statistical significance of a test statistic with an unknown distribution–for instance, the maximum of multiple correlated statistics or some omnibus test statistic for a gene, gene-set or pathway–naive application of permutations may result in an invalid test. The risk of performing an invalid permutation test is particularly acute in complex trait mapping where polygenicity may combine with a structured population resulting from the presence of families, cryptic relatedness, admixture or population stratification. I give both analytical derivations and a conceptual understanding of why typical permutation procedures fail and suggest an alternative permutation based algorithm, MVNpermute, that succeeds. In particular, I examine the case where a linear mixed model is used to analyze a quantitative trait and show that both phenotype and genotype permutations may result in an invalid permutation test. I provide a formula that predicts the amount of inflation of the type 1 error rate depending on the degree of misspecification of the covariance structure of the polygenic effect and the heritability of the trait. I validate this formula by doing simulations, showing that the permutation distribution matches the theoretical expectation, and that my suggested permutation based test obtains the correct null distribution. Finally, I discuss situations where naive permutations of the phenotype or genotype are valid and the applicability of the results to other test statistics.

## Introduction

In the search for genetic determinants of complex traits, we may be faced with the difficulty of determining the statistical significance of a given test statistic which does not necessarily follow any known probability distribution. This arises when correcting for the multiple comparisons of many correlated tests—e.g. to determine genomewide significance [Abney et al., 2002; Cheng and Palmer, 2013]—or in methods where multiple variants (e.g. rare variants) are aggregated into an omnibus test. Methods that use weights that vary depending on the phenotype data, for instance, typically do not have a known asymptotic distribution and require resampling methods to estimate significance [Sha et al., 2012; Fang et al., 2012]. Even when an asymptotic distribution is known, realities of genetic data, such as population structure or linkage disequilibrium, may result in an inflated false positive rate [Tintle et al., 2011; Epstein et al., 2012; Liu et al., 2013]. Family based methods, though often robust to population stratification, also can have false positive rates above the nominal level [Kazma and Bailey, 2011; Greco et al., 2014]. Permutation tests can be a solution in such cases [Basu and Pan, 2011; Lin and Tang, 2011], but rely on the assumption that the subjects are independent. This assumption is violated, for instance, in the presence of population stratification [Epstein et al., 2012; Liu et al., 2013] or familial relatedness [e.g. Abney et al., 2002; Bourgain and Genin, 2005; Kazma and Bailey, 2011] preventing the valid application of a permutation test.

At the heart of the invalidity of a permutation test in the presence of population stratification or relatedness is the presence of polygenic effects and its confounding with genotypes. As I discuss below, this may result in a lack of exchangeability between subjects, a fundamental requirement of a permutation test. It is worth noting that relatedness is not always a barrier to a valid permutation test. For instance, in some model organism breeding designs, exchangeability exists, allowing a valid permutation test [Churchill and Doerge, 1994], and more complicated breeding designs can also, with careful thought, lead to valid permutation tests [Churchill and Doerge, 2008; Peirce et al., 2008; Cheng et al., 2010; Cheng and Palmer, 2013]. Similarly, given specific restrictions on the types of relatedness that is present among the subjects (e.g. only siblings), it may be possible to formulate a valid permutation test[e.g. Allison et al., 1999; Fang et al., 2012]. However, many forms of population structure, including familial relatedness, can cause confounding that can invalidate a permutation test. Although in a simple population stratification scenario—where a limited number of principal components can adjust for the background genetic confounding—it is possible to formulate a valid permutation test [Epstein et al., 2012], in more complicated scenarios, as often exist in human studies—where close or distant relatedness, cryptic or otherwise, may possibly combine with other forms of population structure—a clear statistical framework to help researchers determine the applicability of a permutation test, or how precisely to do such a test, has been lacking.

Here, I consider possible permutation approaches for quantitative traits where arbitrary forms of population structure may exist in the sample. The presence of both population structure and polygenicity leads me to using the linear mixed model (LMM) for multivariate normal data as a foundation on which to build, as this is a standard model used in the genetic analysis of quantitative traits. Although the approaches used here may be applicable to non-normal types of data, I do not consider this issue. Certainly, permutation tests in LMMs have been considered in the past (e.g. [Anderson and Robinson, 2001; Anderson and Ter Braak, 2003]), however these studies consider the case where either the “treatment” (e.g. genotypes) is assigned randomly by the researcher or the stochastic components of the model (i.e. the random effect plus the error terms) are independent. Neither of these situations generally holds true in the genetic analysis of a complex trait, where the researcher is not at liberty to assign genotypes at random and the polygenic effect generally results in non-independence of the random effect. In addition to defining the LMM, I show how misspecification of the covariance matrix leads to an altered asymptotic distribution of the standard test statistic, and how different permutation approaches, can be modeled through different forms of misspecification of the covariance matrix. I discuss this issue further below. Finally, I also discuss what precisely should be permuted, phenotypes, residuals, or genotypes, and provide simulation results supporting the analytical findings.

## Statistical Model

Here I define the statistical model used in the remainder of this article and the resultant likelihood. Given this model, I propose a standard test statistic which, under the right set of conditions, asymptotically follows a central chi-squared distribution with one degree of freedom 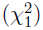. Although it is not really necessary to use a permutation test when the distribution of the statistic is known, its analytical tractability allows for insights that also apply to more general cases. Given the model, I then define exchangeability and the conditions that are needed to ensure that exchangeability holds.

Given *n* subjects with phenotype data **y** = (*y*_1_, …, *y*_*n*_)^*t*^, where the superscript ()^*t*^ indicates transpose, the *n* × *p* covariate data matrix **X**, which includes the intercept term, and the predictor of interest (e.g. genotypes) g = (*g*_1_, …, *g*_*n*_)^*t*^, the LMM is

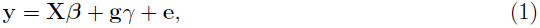

Where *β* is the vector of parameters for the covariates, γ is the scalar parameter for the predictor of interest, and e ~ MVN(0, **∑***σ*^2^) is an error term. The error term encompasses both a random effect and residual error **e**\*, **e** = **u** + **e**\*. The residual errors are distributed as independent normals with variance 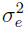. In the genetic context the random effect u will typically be the polygenic effect, and if we further assume that it is the sum of a large number of independent additive genetic effects in an outbred sample, the central limit theorem dictates that u is multivariate normally distributed with correlation matrix **K** [Lange, 1978] with the result extended to the case of inbreeding and dominance variance in Abney et al. [2000]. Although I do not assume a particular structure for ∑, a common parameterization in a genetic LMM is **∑** = **K***h*^2^ + **I**(1–*h*^2^), where **K** is an additive genetic relationship matrix (GRM), I is the identity matrix and *h*^2^ is the narrow-sense heritability. The matrix K may be estimated from available genotype data or determined from a pedigree, in which case it is equal to 2**Φ**, where **Φ** is the matrix of kinship coefficients. The log likelihood given this model is(2)

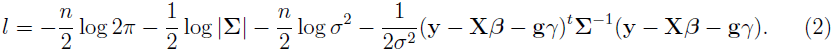

The quantity of interest is the parameter *γ*, and under the null hypothesis *γ* = 0. To test against the alternative *γ* ≠ 0, the statistic 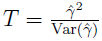 where γ̂ is the best linear unbiased estimator (BLUE) (equivalently the maximum likelihood estimator) of *γ*, has a 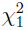 distribution under the null hypothesis when *σ*^2^ is known. In practice, we use

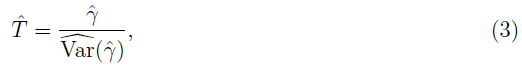

where the estimation Variace 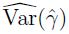 uses an estimator *S*^2^ in place of the true variance *σ*^2^. In an LMM approach 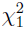 and is an unbiased estimator of *σ*^2^. This results in *T̂* being asymptotically *X*^2^_1_ distributed. However, in a genetic analysis **∑** is not always known, leading to the question of what the distribution of *T̂* is when **∑** is misspecified.

In genetic analyses of complex traits, using an LMM with a misspecified covariance matrix is likely a common occurrence. Perhaps the simplest example of this is when unrelated individuals are unknowingly sampled from two populations, with different allele frequencies at the tested marker, but are assumed to be from a single population. If the trait is associated with population membership, we see an inflated false positive rate. This sort of confounding is easily corrected by including either a covariate with an indicator of population membership or a block structured correlation matrix with elements equal to 1 when a pair is from the same population or 0 when they are not. At the other end of the population structure scale, misspecification may occur in family studies with a known pedigree when the pedigree is where the estimated variance wrong or incomplete. In fact, even if the pedigree is known without error, misspecification exists when the kinship matrix (as computed from the pedigree) is used as the additive GRM because under a polygenic model the kinship coefficients give only the expected identity by descent (IBD) sharing across the genome whereas the correlation in phenotype values will be the result of the realized IBD sharing. In spite of this last form of misfit, the successful use of the kinship coefficient in the GRM over many decades of pedigree studies in both humans and animals suggests a degree of robustness to the use of the expected covariance in place of the realized covariance.

In order to quantify the effects of covariance matrix misspecification on hypothesis testing in the LMM, in the Appendix I derive the distribution of the test statistic *T̂* in the case where the incorrect matrix **Ψ** is used instead of **∑** I find that *T̂* is asymptotically distributed as a scaled chi-squared distribution, 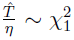 where is a *η* constant. In the case of no covariates and assuming that both y and g have been centered by their mean values, *η* takes the form

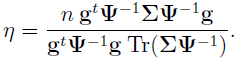

The scalar *η* is, in essence, the genomic control [Devlin and Roeder, 1999; Bacanu et al., 2002] parameter, and its general analytical form in the asymptotic limit of very large sample sizes is in the Appendix. In any real data set we do not know **∑** and, hence, cannot know *η* but having an analytical form will allow us to determine the degree of miscalibration in *T̂* for hypothesized circumstances, as we will see below.

## Confounding, Exchangeability and Permutations

If we expect to perform a permutation test given purely observational data, we should also be concerned with the possibility of confounding. Consider the linear model *y* = *μ* + *x*_1_*β*_1_ +*x*_2_*β*_2_ + *e*, where *e* is an independent error and we wish to test the null hypothesis *β*_2_ = 0. In a designed experiment we can ensure that *x*_2_ has no confounders by random assignment of its values to each subject, and we can safely permute the labels of *x*_2_ to obtain a valid test. With purely observational data, however, *x*_2_ may be confounded with *x*_1_ (due to unknown structure in the data), and permutations of the *x*_2_ subject labels would result in an invalid test. Note that joint permutation of *x*_1_ and *x*_2_ would be valid, but this strategy fails when *x*_1_ is not observed. This situation arises in genetic studies when there is population structure in the sample. In this case *x*_2_ would be genotype and *x*_1_ is a predictor that also depends on the population structure, for example an indicator of population membership (there would be *P* – 1 such indicators for *P* populations in the sample) with each population having a distinct effect on the outcome *y*. If genotype *x*_2_ is dependent on the population structure, we would not want to naively permute all the subject labels of *x*_2_, as this would give an incorrect type 1 error rate. In this case, even if population membership is not recorded, it can often be inferred if there is sufficient genetic data.

Less well appreciated is that, given structured data realized in a covariance matrix, confounding can occur even when the predictor *x*_2_ is independent of the unobserved structured-population predictor *x*_1_. Note that this independence is conditional given the covariance matrices (representing population structure) for *x*_1_ and *x*_2_ in the sense that they are vectors drawn independently from two distinct multivariate distributions each with a given covariance matrix. With observational data, *x*_2_ and *x*_1_ are effectively random vectors, and if *x*_2_ and *x*_1_ have the same, or broadly similar, covariance matrices the outcome *y* will be confounded with *x*_2_, even though *x*_2_ and *x*_1_ are (conditionally) independent. We can gain an intuitive understanding of this by considering an example where the subjects are connected by some pedigree with *x*_2_ being their genotypes at a marker that has no genetic effect and is not in linkage disequilibrium with any causal locus, and with *x*_1_ representing the polygenic effect. The genotype and polygenic effect necessarily have the same covariance matrix **K** induced by the pedigree. The polygenic effect is not observed, so consider the covariance **∑** it induces on the trait *y*. The matrix **∑** is similar to **K**, depending on the heritability. Thus, even though *x*_2_ and *y* are independent, similar genotype values will tend to match-up with similar trait values simply because these random variables have a similar correlation structure. That is, consider some subjects who happen to have high genotype correlation. These subjects will tend to have, for instance, genotype values equal to 2. They will also have high correlation in the trait—assuming a high heritability—and, for instance, will all have low trait values. The net effect is that the 2 genotype will appear to be associated with low trait values, even though these random variables were generated independently. Note that this argument does not depend on whether **K** is the result of population structure resulting from a pedigree or the block diagonal form, with constant off diagonals in each block, that results from assuming a population specific genetic effect. Every genetic trait will depend on genotypes with some population structure covariance **K**, resulting in confounding when testing a genetic marker that also has covariance **K**, thus altering the type 1 error away from the expected amount unless the confounding is corrected for in the test [Newman et al., 2001]. Conversely, if the elements of either *x*_2_ or *x*_1_ are independent—hence, either one has the identity as the covariance matrix—there will be no confounding of *x*_2_ with *y*. For instance, if the *x*_2_ genotypes were independent binomials, there would neither be inflation of the test statistic, nor any problems with permuting the values of *x*_2_. Unfortunately, with observational data verifying the absence of confounding, and the permissibility of a permutation test, may not be possible.

Developing a permutation test for observational data requires assessing whether the permuted quantities are exchangeable. Quantities are exchangeable if, upon permutation of the labels of those quantities, their distribution function is unchanged [Bernardo and Smith, 2000, Sec. 4.2]. In particular, because we want to know the distribution of the test statistic under the null hypothesis, we require exchangeability when *γ* = 0. In an LMM the natural quantities to permute are the residuals **e** = **y** — **Xβ**. In the Appendix I show that the residuals are exchangeable only in the special case where **∑**_*ii*_ = *a* and **∑**_*ij*_ = *b*, *i* ≠ *j* for some scalar constants *a* and *b*, where ∑_*ij*_ is the *i*, *j*-th element of **∑** Note that because we do not in general know *β* or *σ*^2^ but must instead estimate them, even when ∑ has an exchangeable structure permuting the residuals technically provides only an approximate permutation test, though the approximation tends to be very accurate [Anderson and Robinson, 2001].

In general, when using an LMM to model polygenic variation the matrix **∑** will not have an exchangeable structure. Nevertheless, we might undertake a permutation test where the residuals are permuted under the assumption that the phenotype has an exchangeable correlation matrix **Ψ** rather than true correlation matrix **∑**. The fundamental question is will these permutations give an unbiased estimate of the threshold for rejecting the null hypothesis at some specified false positive rate? To address this question exactly, we would need to understand the properties of the order statistics *T*_(*k*)_, *k* = 1, …, *n*! under permutations of the residuals. Instead, I address this in an approximate, but more intuitive, approach by treating the statistics *T*_(*k*)_ as samples from a distribution with covariance matrix that has an exchangeable structure. In the simulation results below we will see that the empiric distribution we get by doing permutations closely matches the distribution obtained from assuming **Ψ** = **I**.

## Simulations

The simulations are done in a sample of 1,415 Hutterite individuals with a known 13 generation pedigree [Abney et al., 2000]. Phenotypes for the sample are generated under the null model to have a mean of 3.0 and covariance matrix **∑***σ*^2^ with **∑** = 2**Φ***h*^2^ + **I**(1 –*h*^2^), with **Φ** the kinship coefficient matrix as computed from the pedigree and *h*^2^ the narrow sense heritability. Genotypes are simulated by randomly assigning the founders of the pedigree a genotype from a biallelic marker with minor allele frequency of 0.3 and using Mendelian segregation to randomly determine the genotypes of all the other pedigree members.

First, I address the question of what happens when “naive” permutations are done. That is, the residuals under the null model are permuted regardless of whether the correlation matrix is exchangeable and the new phenotype (i.e. the covariate effects plus the permuted residuals) is put through the same LMM analysis as the original data. More precisely, we assume the null model **y** = **X*β*** + **e** where **e** ~ MVN(0, *σ*^2^**∑**). That is, analyses done under the null model use exactly the same model as that used to generate the data. Using equation (2) with γ = 0, I first fit the null model and obtain maximum likelihood estimates for the parameters, 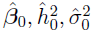 Using generalized least squares (GLS), I test the null hypothesis γ = 0 against the alternative γ ≠ 0 using the test statistic *T̂* (3) computed under the alternative model,

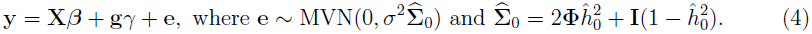

Note that *T̂* is necessarily asymptotically distributed as a 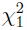 because 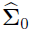 asymptotically converges to the true correlation matrix **∑** Also note that *σ*^2^ in equation (4) is estimated by the sample variance when computing *T̂*. I want to compare this asymptotic distribution with the empirical distribution one obtains by doing naive permutations. To do this I first obtain the estimated residuals under the null, **ê** = **b** – **X*****β***_0_. I permute these residuals to obtain **ê**_1_ and a new phenotype vector **y***π*_1_ = **X***β̂*_0_ + **ê***̂*_1_ Under the alternative model of equation (4) but with **y***̂*_1_ in place of **y**, I obtain a test statistic 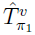, where the *v* superscript indicates the use of naive permutations. I repeat this process *L* = 10^4^ times to obtain 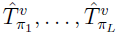 If doing naive permutations were to provide the correct empiric distribution for our original test statistic *T̂*, then the samples 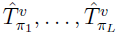 should follow a 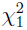 distribution.

As shown in Figure 1 the empiric distribution clearly fails to follow the desired asymptotic distribution. The reason for this is that the permutations fail to maintain the correlation structure of the original phenotype data. As discussed above, this form of permutation would give an accurate distribution only when the true correlation matrix of the estimated residuals has an exchangeable structure. We can use the methods in the Appendix to quantify the inaccuracy of the empiric permutation distribution. We can model the statistics 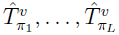 as coming from the distribution 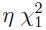 that results from assuming the incorrect correlation matrix **Ψ** = **I** rather than the correlation matrix 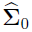. Using the theoretically computed value of *η*, as given in the Appendix, and plotting 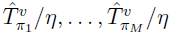 against a 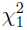 in Figure 1(B) we see that the distributions match well. From this we see that computing the significance of *T̂* from 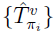 would lead to an anti-conservative estimated level of significance. For instance, to get a nominal levels of significance of 10^−4^ and 10^−5^, the permuted distribution would select threshold levels of 12.7 and 16.3, respectively. Because the actual distribution of the test statistic is 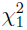, the observed type 1 error rates would be 3.7 × 10^−4^ and 5.4 × 10^−5^, respectively, a substantial inflation.

**Figure 1.**
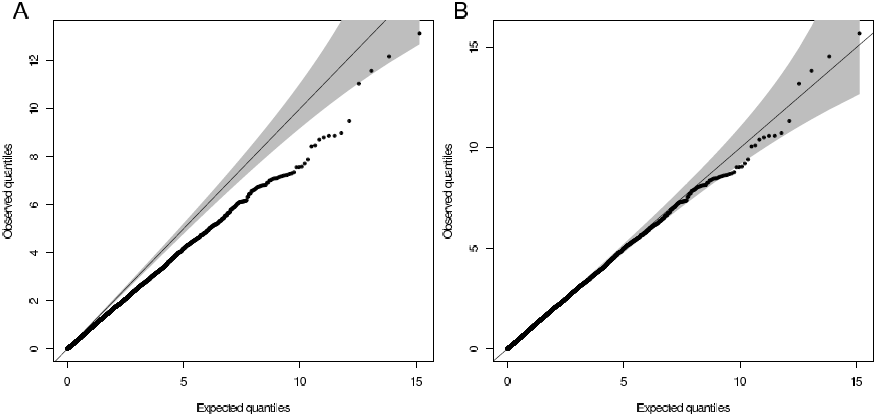
QQ plots under naive phenotype residual permutations. In both plots the expected quantiles are for a 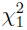 distribution and the shaded area is the 95% confidence region. (A) The observed quantiles are the values of the test statistic under permutations of the trait values. (B) The observed quantiles are the values in (A) divided by the theoretical inflation factor.

Though it is not possible to do a exact permutation test when the residuals have a non-exchangeable correlation matrix, it is possible to do a valid permutation-based test. The approach (referred to here as MVNpermute) is described in Abney et al. [2002] and the Appendix and relies on the fact that there exists a linear transformation of the residuals that results in a vector (i.e. the transformed residuals) whose covariance matrix is proportional to the identity matrix, and is therefore exchangeable. Because MVNpermute is based on permutations of an invertible transformation of the phenotype residuals, all structure in the genotype data (e.g., linkage disequilibrium, allele frequencies) is preserved. Inverting the transformation following permutations then results in new simulated data sets that maintain the structure in the entire original data (i.e., phenotype correlations and genotype structure).

I repeated the above simulations but with the permutations generated using MVNpermute. This gave statistics 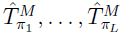 where the *M* superscript indicates the use of MVNpermute. As shown in Figure 2, this results in statistics that follow the expected distribution. That is, by first decorrelating the residuals—ensuring exchangeability for a normally distributed trait—permutations allow us to estimate the proper threshold for a given false positive rate. In practice, MVNpermute is not necessary to determine the p-value at a single SNP, but obtaining *L* permutation-based data sets {**y**_*π_i_*_} allows us to do an empiric multiple testing correction to determine genomewide significance, for instance [Abney et al., 2002].

**Figure 2.**
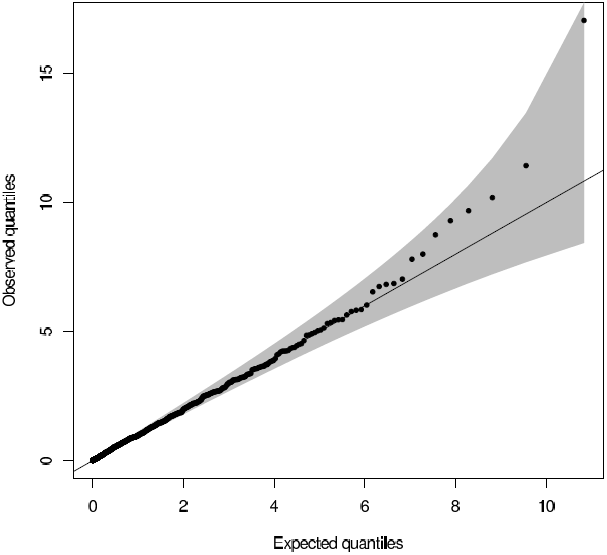
QQ plot of the MVN permute method. The observed quantiles are the values of the test statitic from 10,000 MVN permutations, while the theoretical quantiles are those from a 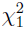 distribution. The shaded region is the 95% confidence bounds.

It is not unusual to realize that permuting the phenotypes (or rather the residuals) does not result in a valid permutation test when individuals are related. A possible alternative is to permute the genotypes instead of the phenotypes. A rationale is that the LMM inference is based on the conditional distribution of the phenotypes given the genotypes. As the phenotypes remain fixed while new genotypes get assigned to individuals via permutation, the correlation structure in the phenotypes is preserved, resulting in a valid permutation test. In fact, many statistics used in complex trait mapping assume a distribution that is conditional on the genotype data, even if the form of the test statistic distribution is not known. Permuting the genotype data, then, to estimate this distribution seems a natural approach.

To understand the consequences of a genotype permutation procedure on an arbitrary test statistic, let us first consider the standard ordinary least squares (OLS) statistic,

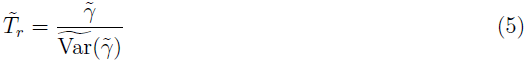

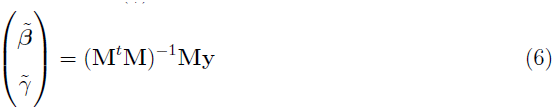

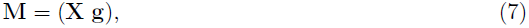

where ·̂ indicates estimation with a scaled covariance matrix **Ψ** rather than **∑** (in this case **Ψ** = **I**). That is, the statistic we use does not explicitly account for the polygenic effect. We may be aware that relatedness in our sample will result in *T̂*_*r*_ not being 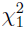 distributed and, thus, perform genotype permutations to obtain the empiric distribution of *T̂*_*r*_. To see if genotype permutations recover the correct distribution, I performed *L* = 10^4^ permutations of the genotype data while keeping the phenotype data constant to obtain {*T̂*_*r*,*π*_*1*__,…, *T̂*_*r*,*π*_*L*__} where *π*_1_, …., *π*_*L*_ index the permutations. I then compare this to a sample from the true null distribution that I obtain by performing *L* gene dropping simulations, *T̂*_*r*,1_, …, *T̂*_*r*,*L*_. The results in Figure 3 show that the distribution obtained by genotype permutations is highly deflated relative to the distribution from gene dropping. Using this approach to obtain an empiric threshold of significance would result in a highly inflated false positive rate.

**Figure 3.**
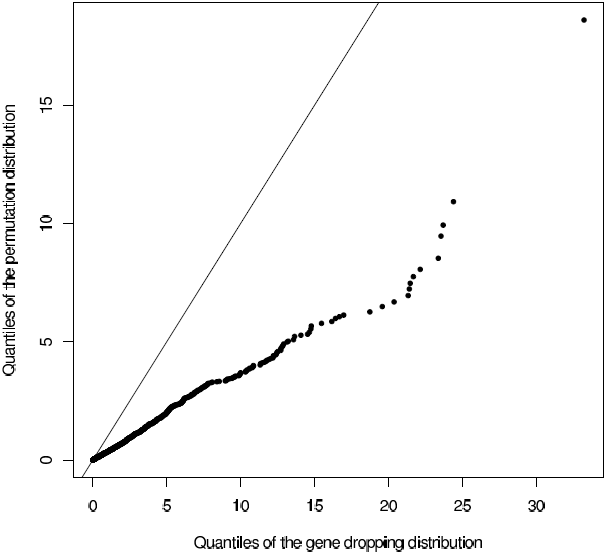
QQ plot of the empiric null distribution for the ordinary least squares statistic against the expected null distribution. The expected null distribution is a sample obtained by doing gene dropping. The solid line is the *y* = *x* line.

The source of the problem with genotype permutations can be understood by returning to the notion of confounding. Because genotypes were not randomly assigned by the researcher to subjects, the absence of confounding is not guaranteed. In fact, the subject genotypes are correlated as a consequence of Mendelian segregation and all markers in the genome, whether causative or not, share the same pedigree for a given set of individuals. Hence, the covariance of the marker being tested is equal (up to a scalar constant) to the covariance of the polygenic effect. If genotypes are permuted, the similarity of the covariance structures of phenotype and genotype will not be preserved. Thus, I expect that the null distribution of an arbitrary test statistic, not just the OLS statistic, will not be correctly estimated by genotype permutations, in general.

Although the null distribution of an arbitrary test statistic cannot be inferred from genotype permutations, the null distribution of the GLS statistic can be. That is, if instead of using *T̂*_*r*_ we use *T̂* as defined in equation (3) and do the genotype permutation procedure as described above, we find the permutation distribution of *T̂* matches the gene dropping permutation distribution (data not shown). We can understand this result by looking at the definition of the GLS statistic *T̂*. We can view the GLS statistic as the OLS statistic computed on the data following a decorrelation step. That is, if we define **∑**^1/2^ as the symmetric square root matrix of **∑** and **z** = **∑**^−1/2^**y**, **W** = **∑**^−1/2^**X**, **f** = **∑**^−1/2^**g**, *ϵ* = **∑**^−1/2^**e**, then we obtain the linear model **z** = **W*β*** + **f**_γ_ + ***ϵ***, with ***ϵ*** ~ MVN(0, **I**σ^2^). The GLS statistic on the original data is equivalent to the OLS statistic on the decorrelated data z. Because our new trait data **z** are normally distributed and uncorrelated, they are independent and can no longer be confounded with the genotypes under the null hypothesis. In the absence of confounding, then, permuting the genotypes recovers the true null distribution of the test statistic *T̂*.

## Software

The MVNpermute algorithm is implemented in the R programming language and is available for download from the Comprehensive R Archive Network (http:\\cran.r-project.org) as the “MVNpermute” package.

## Discussion

The fundamental challenge with performing a permutation test is ensuring exchangeability in the permuted quantities. In a genetic association test it is generally not possible to do an exact permutation test when the trait under study has a polygenic component. The reason is that confounding due to population structure exists between the genotype being tested and the unknown polygenic effect, both of which have similar covariance structures. Only when all individuals are equally related, as in an F2 cross [Churchill and Doerge, 1994], will a naive permutation approach obtain the correct type 1 error rate. Nevertheless, with an accurate estimate of the trait covariance structure, it may be possible to remove the confounding and perform a valid permutation test. I have described an approach we have previously proposed [Abney et al., 2002] for removing the correlation in the phenotype residuals and shown that it generates the correct null distribution. Strictly, the method is valid when the phenotype data are multivariate normally distributed, where removing the correlation is sufficient to ensure exchangeability. Another permutation approach was proposed by Aulchenko et al. [2007]. They estimate the polygenic effect and obtain estimates of the residual error term. Although, under multivariate normality, the residual errors in equation (1) are exchangeable, the *estimated* residual errors, in general, will not be. Nevertheless, this may be a case of “close enough,” allowing for a reasonably accurate estimate of significance thresholds, though I have not investigated this question.

Other resampling strategies are possible, though they also have limitations. Gene dropping is one such approach. In this strategy one simulates the Mendelian segregation of the founder genotypes through all descendents. Because Mendelian segregation is random and independent of the phenotype, it provides a valid distribution of the test statistic under the null hypothesis. The primary difficulties with gene dropping is the need for a complete pedigree and knowledge of the founder genotypes. If the pedigree is known, but the founder genotypes are not, it may be possible to reconstruct, or simply guess, them from the available data. Doing so, however, runs the risk of introducing unknown biases as the observed genotypes may be confounded with the phenotypes. On the other hand, if the pedigree is not known, gene dropping is simply not feasible.

Instead of gene dropping we might try to permute genotypes, leaving the covariance structure of the phenotype intact. As discussed above, for an arbitrary test statistic this does not necessarily result in a valid test as permutations of the genotypes will not preserve their covariance structure. In addition, applying a “decorrelating” transformation to the genotypes is not sufficient to ensure their exchangeability because, unlike the multivariate normal distribution of the phenotype, the joint distribution of the genotypes has higher order dependencies. Nevertheless, in the case of a multivariate normal phenotype being analyzed with a linear mixed model, the standard test statistic *T̂* naturally transforms the trait data to being independent. Once independent, any sets of dependent or independent genotypes, including permutations of the original ones, can be used to recover the correct null distribution of the test statistic. This approach has been used in mouse cross data to obtain proper genomewide significance levels [Cheng et al., 2010; Cheng and Palmer, 2013], and more recently in humans [Zhang et al., 2014]. It seems likely that any test statistic that removes the correlation in the phenotype data and does not depend on Mendelian segregation under the null hypothesis, would allow genotype permutations to be valid, though I have not investigated this further. An additional caveat arises, however, with genotype permutations when there are other covariates. In particular, if any of the covariates are associated with the genotype, genotype permutations may estimate the incorrect null distribution. That is, it will give the null distribution for when the covariate and genotype are not associated rather than for when they are. This may arise, for instance, when a covariate is itself a genetic trait, or when it is a principal component vector obtained from a population structure analysis. It may also arise when testing effects such as gene by environment interaction. In this situation the null model has a nonzero genetic main e ect. In general, an approximate permutation test of interaction effects is done by computing and permuting outcome residuals [Anderson and Ter Braak, 2003] as done by MVNpermute. Additional work is needed to understand this effectiveness and validity of MVNpermute and genotype permutations in the presence of genetic interaction terms and of genotype permutations when other genetic predictors are in the null model.

Another resampling strategy is the parametric bootstrap. In this approach one assumes the phenotypes follow a particular parametric distribution with parameter values equal to those estimated from the observed data under the null hypothesis. Samples are then drawn from this distribution and a test statistic computed for each sample, thus obtaining an empiric null distribution. For instance, one might assume the phenotype follows a multivariate normal distribution with fixed effect parameters and variance components estimated by maximum likelihood under the null model. Drawing many phenotypes from this distribution and testing the genotype at a SNP against each randomly drawn phenotype provides a null distribution for the test statistic. This approach relies on the parametric distribution accurately representing the observed data. Insofar as the data deviate from the assumed distribution, biases in the estimated significance threshold may ensue. A true permutation test has the advantage of not needing to make such parametric assumptions. The MVNpermute method also relies on certain distributional assumptions. Namely, that exchangeability under the null is determined by the structure of the correlation matrix. Intuitively, this assumption appears weaker than those used in a parametric bootstrap, suggesting that there may be greater robustness to the permutation based approach, though I have not investigated this question.

The analyses I performed here were based on using a statistic known to asymptotically follow a 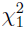 distribution. This allowed me to easily show the invalidity of particular permutation procedures. In practice, one would not need a permutation test for such a statistic, but the lessons extend to other statistics as well. For instance, we might want to determine statistical significance after correcting for multiple correlated tests, as when doing a genomewide scan or a scan over a smaller region. In this case the statistic of interest would be the maximum over all statistics in the scan. Similarly, statistics that jointly combine information across SNPs or use phenotype dependent weights, for which there may not be a clear generative model, may not have a known distribution under the null hypothesis. Situations such as these would benefit from a permutation test, if one exists. The presence of polygenic variation may make a true permutation test difficult or impossible, but a permutation based test may be achievable by carefully considering the sources of correlation, or non-exchangeability, in the data. Hopefully, the examples and discussion I provided here will help bring insight into the development of such tests.

## Acknowledgements

I would like to thank Abraham Palmer, Riyan Cheng, Peter Carbonetto and Lei Sun for repeatedly raising this topic with me and motivating me to write this manuscript. Peter Carbonetto also provided helpful comments on the MVNpermute R code. This work was supported by NIH grant HG002899.

The author has no conflicts of interest to declare.

## Appendix

### Distribution with a misspecified covariance matrix

The LMM of the main text is **y** = **X*β*** + **g**γ + e, where e ~ *N*(0,**∑***σ*^2^). The asymptotic distribution of the test statistic 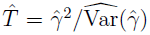 is 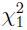 under the null hypothesis, but only if we use the correct scaled covariance matrix **∑** in our estimate of γ̂ and its variance. If we misspecify this matrix, the distribution of *T̂* becomes a scaled chi-squared distribution 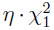 with the scalar *η* depending on the amount of misspecification. To determine *η* we can derive *T̂* and its distribution assuming that we have used the incorrect covariance matrix **Ψ***σ*^2^ in place of the correct **∑***σ*^2^. First, we obtain the BLUE for γ. If we let **Ψ**^1/2^ be the symmetric positive definite square root matrix of **Ψ** and define

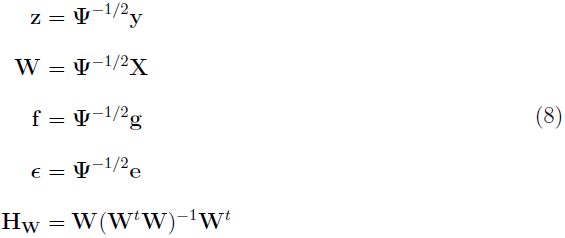

then the BLUE for γ is

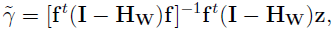

where ·̂ indicates an estimate using **Ψ** rather than **∑**. The variance of this estimator is

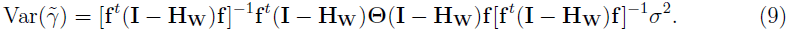

where Var(z) = **Θ***σ*^2^ = **Ψ**^−1/2^**∑Ψ**^−1/2^*σ*^2^. The statistic

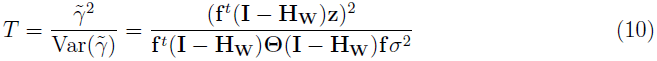

is 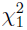 because z is multivariate normal. The statistic *T*, however, is not an adequate test statistic because it depends on the unknown matrix **∑** and the unknown parameter *σ*^2^. In practice we use the test statistic

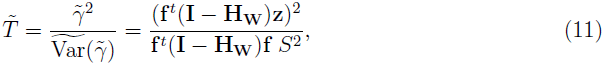

where in place of *σ*^2^ we have the sample variance

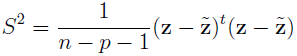

where ẑ = **H**_**A**_**z** = **A**(**A**^*t*^**A**)^−1^**A**^*t*^**z** and **A** = (**W f**) = **Ψ**^−1/2^**M**, **M** = (**X g**). It is straightforward to show that if **Ψ** = **∑** then the expectation *E*(*S*^2^) = *σ*^2^. In general, however, we have,

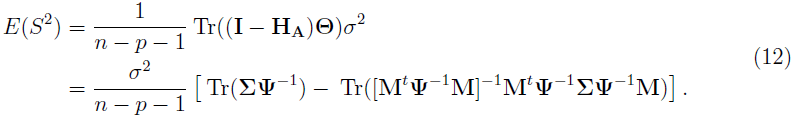

The result of misspecifying the covariance matrix is given by the following lemma.

#### Lemma 1.

Let y ~ *N*(**X***β* + gγ, **∑***σ*^2^) with **∑** non-negative definite and **Ψ** be some symmetric non-negative definite matrix. Define **Θ** = **Ψ**^−1/2^**∑Ψ**^−1/2^, **M** = (**X g**), with **z, f, H**_**W**_, **Θ** as in equation (8) and **T̂** as in equation (11). If *λ*_1_(**∑Ψ**^−1^) = *o*(*n*^1/2^), where λ_1_(**∑Ψ**^−1^) is the largest eigenvalue of matrix **∑Ψ**^−1^ (equivalently the largest eigenvalue of **Θ**), then as *n* → *∞*

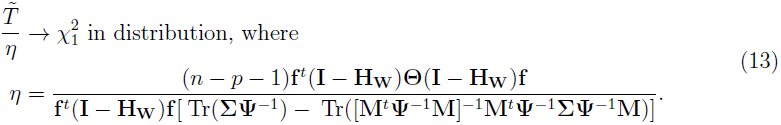

*Proof* We can write 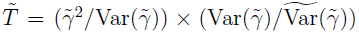, which is the product of a 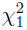 random variable, as given in equation (10), and the ratio of the true to estimated variance of γ̂. The true variance is given by equation (9), but the estimated variance is

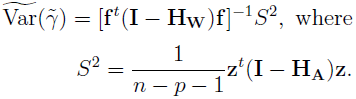

Thus, the test statistic is

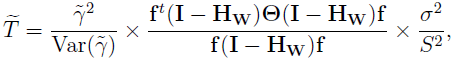

the product of a 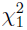 random variable and the quantity

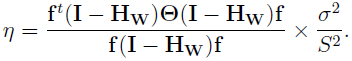

If as *n* → ∞ Var(*S*^2^) → 0, then 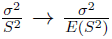 in probability, where *E*(*S*^2^) is given by equation (12). Thus, we obtain equation (13) of Lemma 1 when Var(*S*^2^) → 0. We can obtain a su cient condition for Var(*S*^2^) → 0 by considering

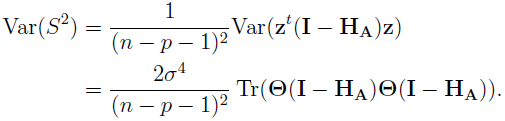

Thus, Var(*S*^2^) → 0 when Tr(**Θ**(**I** – **H**_**A**_)**Θ**(**I** – **H**_**A**_)) = *o*(*n*^2^).

Now, consider the following eigenvalue result [Zhang, 2011, Theorem 8.12, p. 274]. Let λ_*i*_(**P**) be the *i*th eigenvalue for some *n* × *n* matrix **P** ordered such that λ_1_(**P**) ≥ λ_2_(**P**) ≥ ··· ≥ λ_*n*_(**P**). Then, for any *n* × *n* non-negative definite, Hermitian matrices **P**, **Q**

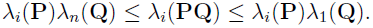

It follows that λ_*i*_(**Θ**[**I** – **H**_**A**_]) ≤ λ_*i*_(**Θ**) because **Θ** is symmetric, non-negative definite and **I** – **H**_**A**_ is symmetric and idempotent with all eigenvalues equal to 0 or 1. Furthermore, **Θ**[**I** – **H**_**A**_] is non-negative definite because λ_*i*_(**Θ**[**I** – **H**_**A**_]) ≥ λ_*i*_(**Θ**) λ_*n*_(**I** – **H**_**A**_) 0, and thus λ_*i*_(**Θ**[**I** – **H**_**A**_])^2^ ≤ λ_*i*_(**Θ**)^2^. Recalling that the trace of a matrix is the sum of the eigenvalues we have Tr[**Θ**(**I** – **H**_**A**_)(**I** – **H**_**A**_)] ≤ Tr[**Θ**^2^]. If we define **B** = **∑Ψ**;^−1^, then Tr(**Θ**^2^) = Tr(**B**^2^).

Hence,

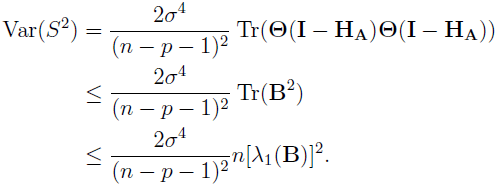

Thus, a sufficient condition for Var(*S*^2^) → 0, and hence *σ*^2^/*S*^2^ → *σ*^2^/*E*(*S*^2^) in probability, is *λ*_1_(**∑Ψ**^−1^) = *o*(*n*^1/2^).

Note that a possibly tighter sufficient condition is 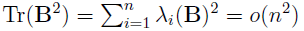

### Exchangeability of the multivariate normal distribution

Given a random vector **y** = (*y*_*1*_, …,*y*_*n*_) distributed as a multivariate normal *f*(**y**) = *N*(*μ*, **∑**), under what conditions are the elements of **y** exchangeable? If we let **P** be a permutation matrix so that **Py** is a permutation of the elements of **y**, then **y** is exchangeable when *f*(**y**) = *f*(**Py**) for every permutation matrix **P** [Bernardo and Smith, 2000, Sec. 4.2]. Taking the log of both sides, this reduces to,

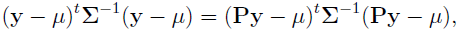

which implies the condition,

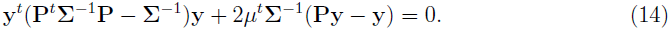

In order for equation (14) to hold for every **y** and **P**, each of the two terms must be zero. If the first term is to be zero for all **y** and **P**, then **P**^*t*^ **∑**^−1^**P** = **∑**^−1^. This condition is met if and only if all the diagonal elements **∑**_*ii*_ = *v*, for some constant *v*, and all the off-diagonals ∑_*ij*_ = *vρ*, for some constant *ρ*. To set the second term to zero we first note that for some vector **w**, the condition **w**^*t*^(**Py** – **y**) = 0 implies **w** = *α*1, for some constant *α* and where **1** = (1,…,1)^*t*^. Hence, given the structure of **∑** we already determined, the second term is zero whenever *μ* = (*μ*,…, *μ*)^*t*^ for some constant *μ*.

In the text, the null model corresponds to **y** ~ *N*(*Xβ*,**∑**). Let us assume the **∑** has an exchangeable structure. In general, however, the vector **X***β* ≠ *μ*1 for some fixed *μ* and y is still not exchangeable. The vector e = **y** – **X*β***, though, does satisfy the requirements for exchangeability, and a permutation test can be based on permutations of the residuals. In practice, the vector *β* is unknown and must be estimated, resulting in estimated residualsê. Permuting the estimated residuals, then, results in only an asymptotically exact permutation test when **∑** has an exchangeable structure.

### MVNpermute algorithm

The permutation based algorithm was originally presented in Abney et al. [2002, pp926–927] and I review it here for completeness. Assume the outcome y follows the model as given in equation (1) in the main text, and let the errors e have known covariance matrix **Ω** = **∑***σ*^2^. In practice, this matrix may not be known, in which case a consistent estimator will maintain the asymptotic properties of the permutation based procedure. For instance, a maximum likelihood estimate under the null model (γ = 0) could be used. The residuals under the null model **ê**_0_ = **y** – **X**β̂_0_, where *β*_0_ = (**X**^*t*^ **Ω**^−1^**X**)^−1^**X**^*t*^ **Ω**^−1^**y**, have covariance matrix **V**\* = **Ω**–**X**(**X**^*t*^**Ω**^−1^**X**)^−1^**X**^*t*^.

The goal, then, is to transform the residuals, which are not exchangeable, to a new vector whose elements are exchangeable. We can accomplish this by pre-multiplying equation (1) by **C**^−*t*^ where **C** is given by the Cholesky decomposition **Ω** = **C**^*t*^**C**. The resulting model under the null hypothesis *γ* = 0 is **z** = **W*β*** + *ϵ* where **z** = **C**^−*t*^**y**, **W** = **C**^−*t*^**X**, and *ϵ* = **C**^−*t*^e. The covariance matrix of the residuals *ϵ̂* = **z** *–* **Wβ̂**_0_ is **V** = **I** – **W**(**W**^*t*^**W**)^−1^W^*t*^. Note that **V** is symmetric and idempotent (i.e., **V**^2^ = **V**) and if **X** is of rank *p* then **V** has rank *n* – *p*. By the spectral theorem we can make the decomposition **V** = **U∧U**^*t*^, where ∧ is a diagonal matrix with the first *n* – *p* elements equal to the eigenvalue 1 and the last *p* elements equal to the eigenvalue 0, and **U** is the matrix whose columns are eigenvectors. Let **U** = (**U**_1_ **U**_0_), where **U**_1_ is the matrix whose *n* – *p* columns are the eigenvectors associated with eigenvalue 1. Then, we have 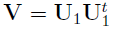 and 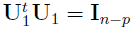. The vector 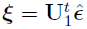 has covariance matrix 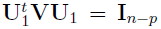 and its elements, under the assumption of multivariate normality of the residuals, are exchangeable. The elements of ***ξ*** are now permuted to obtain ***ξ***^*π*^ = **P*****ξ*** whereP is a permutation matrix, and then transformed by **U**_1_ to get *ϵ̂*_*π*_ = **U**_1_***ξ***^*π*^. Note that I use the convention that “*π*” used as a superscript denotes the variable is permuted, whereas “*π*” used as a subscript denotes that the variable is derived from permuted and non-permuted quantities. A new shuffled data set obtained from the permutation is

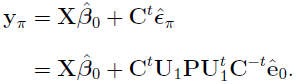

The MVNpermute algorithm is coded as an R function that takes as input the outcome vector y, matrix of covariates **X**, assumed covariance matrix **Ω** and the desired number of permutations. The output is a matrix with columns being the permutation-based outcome vectors. The MVNpermute function is available as a download from CRAN.

